# Vestibular Modulation by Stimulant Derivatives in a Pentameric Ligand-Gated Ion Channel

**DOI:** 10.1101/2024.05.02.592243

**Authors:** Emelia Karlsson, Olivia Andén, Chen Fan, Zaineb Fourati, Ahmed Haouz, Yuxuan Zhuang, Rebecca J Howard, Marc Delarue, Erik Lindahl

**Affiliations:** Department of Biochemistry and Biophysics, Science for Life Laboratory, Stockholm University, Solna, SE-17121, Sweden; Department of Applied Physics, Science for Life Laboratory, KTH Royal Institute of Technology, Solna, SE-17121, Sweden; Cibles Thérapeutiques et Conception de Médicaments (CiTCoM), CNRS UMR 8038, Université Paris Cité, Paris FR-75006, France; Plateforme de Cristallographie et de Cristallogenese, Institut Pasteur, CNRS UMR 3528, Université Paris-Cité, Paris, FR-75015, France; Architecture and Dynamics of Biological Macromolecules, Institut Pasteur, CNRS UMR 3528, Université Paris-Cité, Paris, FR-75015, France

**Keywords:** sTeLIC, allosteric modulation, druggable cavity, ligand-gated ion channel, electrophysiology, cryo-EM, X-ray crystallography

## Abstract

Allosteric modulation of pentameric ligand-gated ion channels (pLGICs) is critical to the action of neurotransmitters and many psychoactive drugs. However, details of their modulatory mechanisms remain unclear, especially beyond the orthosteric neurotransmitter-binding sites. The recently reported prokaryotic channel sTeLIC, a pH-gated homolog of eukaryotic receptors in the pLGIC family, is thought to be modulated by aromatic compounds via a relatively uncharacterized modulatory site in the extracellular vestibule. Here, we show that sTeLIC is sensitive to potentiation by psychostimulant derivatives. By determining new cryo-EM and X-ray structures in closed and open states, and testing the impact of targeted mutations on electrophysiological behavior, we show that several amphiphilic compounds preferentially bind a vestibular pocket in the contracted open-state extracellular domain. This work provides a detailed structure-function mechanism for allosteric potentiation via a noncanonical ligand site, with potential conservation in eukaryotic pentameric ligand-gated ion channels.

## 1 Introduction

Pentameric ligand-gated ion channels (pLGICs) are a widely expressed family of membrane proteins that critically mediate electrochemical signal transduction across evolution [1]. In the animal nervous system, the chemical stimulus of neurotransmitters induces conformational changes, allowing selected ions to pass through the membrane-spanning pore [2]. Members of the pLGIC family are targets in the treatment of neurological and gastrointestinal disorders including epilepsy [3], sleep disruption [4], Alzheimer’s and Parkinson’s diseases [5], nausea and irritable bowel syndrome [6]. Notably, eukaryotic pLGICs share structural architecture and complex pharmacology with prokaryotic channels [7], including conserved sensitivity to clinical drugs such as general anesthetics [8], alcohols [9] and benzodiazepines [10]. Consequently, prokaryotic pLGICs constitute valuable model systems for structure-function studies of gating and modulation throughout the family [11–13].

The minimally conserved topology of each subunit includes an extracellular domain (ECD) comprising ten β-strands (β1 to β10) and a transmembrane domain (TMD) composed of four α-helices (M1 to M4) [12]. Known sites of modulation are also conserved, including so-called orthosteric sites at extracellular subunit interfaces, allosteric modulatory sites facing the outer or inner leaflets of the lipid bilayer, and blocking sites in the ion pore (Figures 1A and 1B) [1]. Recently, small molecules have also been observed to bind in a vestibular site on the internal face of a single-subunit ECD, such as the benzodiazepine flurazepam in ELIC [10] and acetate in GLIC [14].

**Figure 1.**
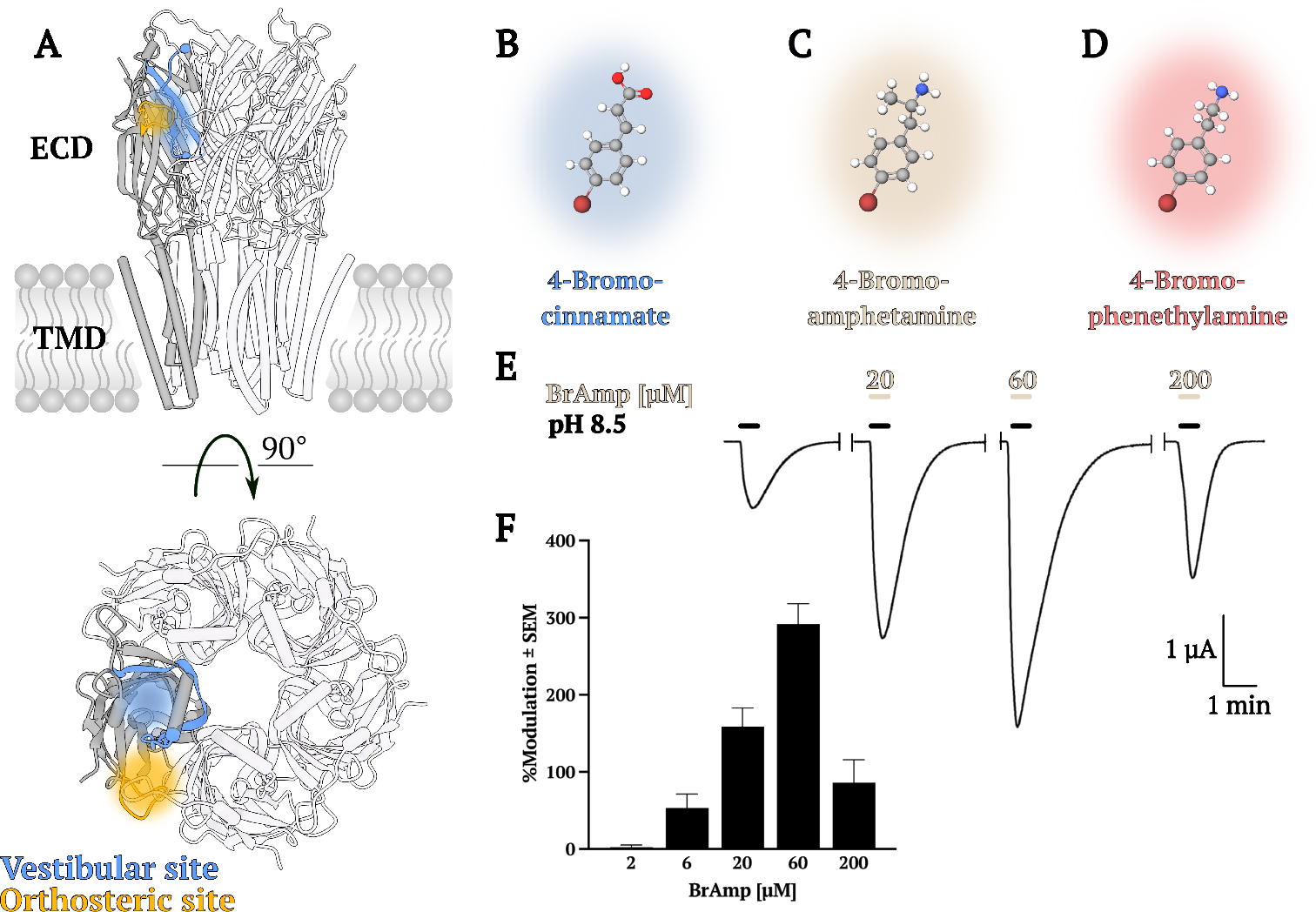
Functional enhancement of a model pLGIC by stimulant derivatives. **A)** Views from the membrane plane (top) and extracellular side (bottom) of a previously published sTeLIC X-ray structure (PDB ID 6FL9) [15], highlighting orthosteric (orange) and vestibular (blue) sites known to bind ligands in at least some pLGIC types. **B)** Ball-and-stick representation of BrCA, a previously reported potentiator binding at the vestibular site. **C)** Model as in (B) of putative modulator BrAmp. **D)** Model as in (B) of putative modulator BrPEA. **E)** Representative electrophysiology recording from an oocyte expressing WT sTeLIC, showing potentiating effects of 30-s coapplications of increasing concentrations of BrAmp with pH 8.5 buffer. **F)** Column plot of sTeLIC potentiation by various concentrations of BrAmp, showing mean percent modulation ± SEM.

A bacterial receptor known as sTeLIC was recently shown to be functionally sensitive to alkaline pH and calcium [15], similar to several eukaryotic receptors [16–18]. sTeLIC was crystallized in a wide-open state with a small synthetic potentiator in the vestibular site (Figures 1A and 1B), which shares structural features with some eukaryotic pLGICs [15, 19]. Based on these observations, we asked whether other drugs might bind in the vestibular site, and how they might modulate function.

Here, we answer these questions using a combination of oocyte electrophysiology, cryo-EM and X-ray crystallography experiments. We show that sTeLIC is sensitive to modulation by amphiphilic compounds including derivatives of amphetamine and phenethylamine psychostimulants. We then leverage both the flexible solvation conditions of cryo-EM and anomalous diffraction available in X-ray crystallography to solve structures in both closed and open states, in lipid nanodiscs and detergent micelles, and in the absence and presence of modulators. From these data, combined with functional characterization of engineered mutants, we find that ligands in the sTeLIC vestibular site preferentially stabilize the open state of the channel, likely by strengthening intersubunit interactions in the ECD. Our results support a distinctive mechanism of allosteric potentiation via the ECD vestibule, apparently specific to a subset of pLGICs including mammalian serotonin type-3 receptors (5-HT_3*A*_Rs).

## 2. Results

### 2.1 Functional Enhancement of a Model pLGIC by Stimulant Derivatives

To investigate the activity of pharmaceutically relevant compounds via the vestibular site in a model pLGIC, we first tested the effect of 4-bromoamphetamine (BrAmp) on sTeLIC in *Xenopus laevis* oocytes. This compound is similar to psychostimulants in common therapeutic and recreational use, with the inclusion of a bromine atom to facilitate structure determination by anomalous X-ray diffraction. We hypothesized that BrAmp would positively modulate sTeLIC, based in part on its structural similarity to 4-bromocinnamate (BrCA) (Figures 1B and 1C) [15]. In two-electrode voltage clamp (TEVC) electrophysiology experiments, BrAmp showed positive modulatory effects similar to BrCA, albeit decreasing in potency at the highest concentrations tested (Figures 1E and 1F). Interestingly, we observed similar potentiation profiles for brominated and unbrominated forms of phenethylamine (PEA) (Figure 1D, Supplementary Figures S1A and S1B), a naturally occurring psychostimulant in bacteria, plants, and animals, including in the human brain [20, 21].

### 2.2 Structures in Lipid Nanodiscs Reveal a Resting State

Given that all previous sTeLIC structures have been reported by X-ray crystallography in an open state [15], we sought a more complete view of the gating cycle by determining the structure in a closed state. Since crystallization conditions were previously shown to favor activation, we pursued single-particle cryo-EM under resting conditions (pH 7.5). To further mimic a physiological environment, we reconstituted the receptor in asolectin with saposin nanodiscs. Interestingly, our initial micrographs nonetheless yielded a single reconstruction with an evidently open pore (Supplementary Table S1, Supplementary Figures S2 and S3). Moreover, we found a distinct non-protein density in the vestibular pocket (Supplementary Figure S1C). This density could reasonably accommodate fluorinated fos-choline-8 (FFC-8), which we had included in grid preparation to combat preferential particle orientation. Indeed, we subsequently found that micromolar concentrations of FFC-8 potentiated sTeLIC currents in oocytes (Supplementary Figure S1C), consistent with binding in this site favoring the open state. This complex was otherwise comparable to other open structures (Cα RMSD *<* 1.4 Å versus PDB ID 6FLI), verifying that sTeLIC conformation is not substantially influenced by X-ray versus cryo-EM methods, nor by detergent versus nanodisc reconstitution.

To minimize channel activation, we then substituted 3-((3-cholamidopropyl) dimethylammonio)-1-propanesulfonate (CHAPS) for FFC-8 during grid preparation. This approach resulted in a C5-symmetric map at a nominal resolution of 2.4 Å (Supplementary Table S1, Supplementary Figures S2 and S3). The associated structure (Figures 2A and 2B) was notably distinct from previous open structures of sTeLIC, with an overall Ca root-mean-square deviation (RMSD) *>* 4.3 Å (versus PDB ID 6FLI). Instead, the pore was comparable to apparent resting structures of the related channel ELIC [22, 23], with a minimal radius too narrow (*<* 2Å) to conduct ions (Figures 2C and 2D). Local resolution varied between extracellular and intracellular ends, with relatively poor definition in the outward-facing half of the ECD (Supplementary Figure S2), reminiscent of the relative flexibility observed in the ECD of the resting-state bacterial channel GLIC [24].

**Figure 2.**
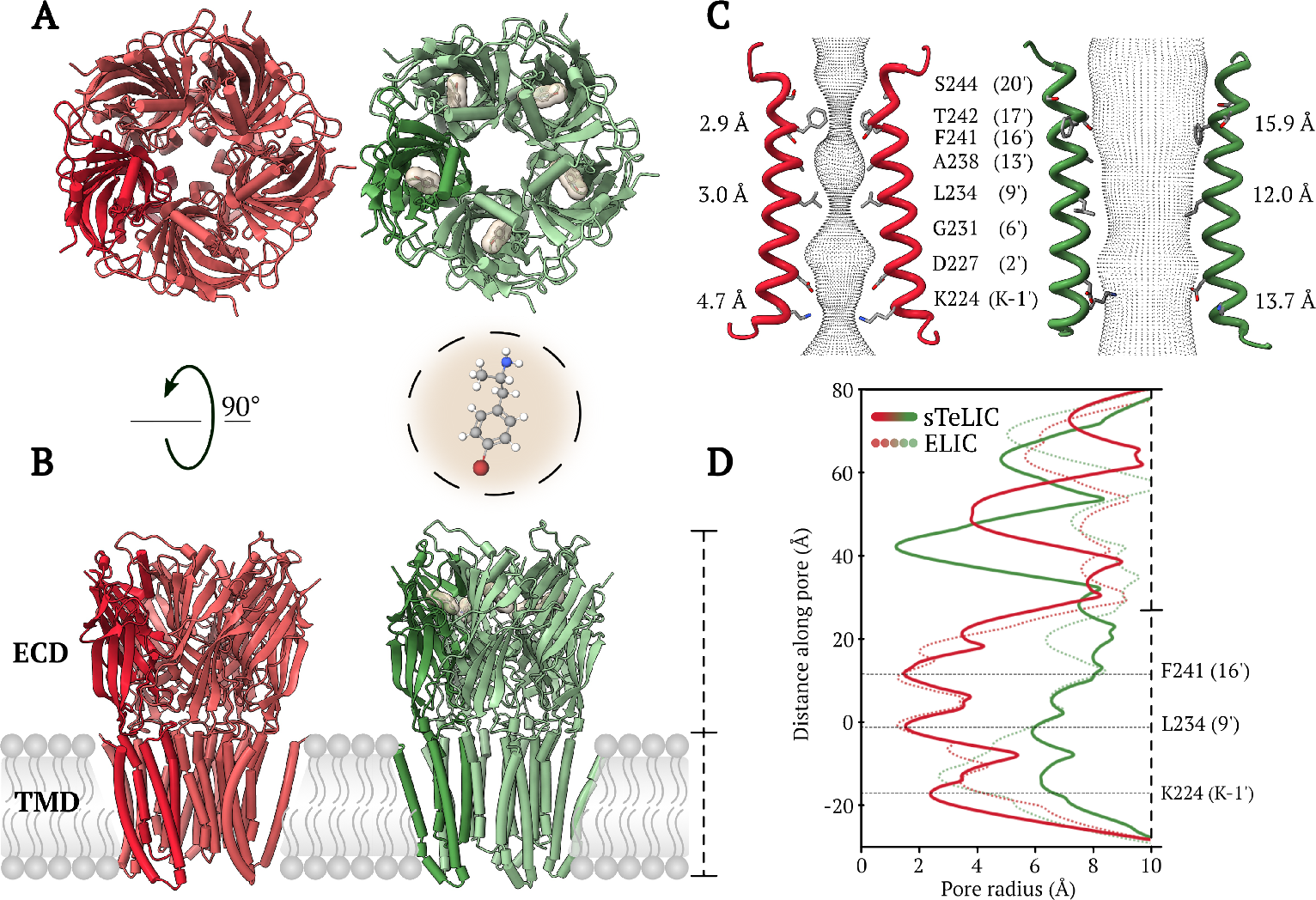
Structures of closed and open sTeLIC in the absence and presence of modulators. **A)** Extracellular and **B)** membrane-plane views of sTeLIC in closed (red) and open (with BrAmp, X-ray, green) states. In each model, one subunit is shaded for clarity. In the open structure, BrAmp is represented as sticks in its molecular surface (beige). In (B), dashed line at right estimates axial span of the channel. **C)** Pore surfaces (gray mesh) calculated using CHAP [26] for the TMD region. For clarity, only two opposing M2-helices are shown, with pore-facing residues labeled (absolute residue positions and prime notation) and shown as sticks. Pore diameters at the -1’, 9’, and 16’ constriction points are indicated beside each model **D)** Pore radius profiles, plotted as a function of distance along the fivefold symmetry axis for closed sTeLIC (solid red), open sTeLIC (with BrAmp, X-ray, solid green), closed ELIC (PDB ID 3RQU [23], dotted red), and open ELIC (PDB ID 8D68 [25], dotted green). Dashed line at right corresponds to axial span shown in (B). Relative to closed structures, the open structures contain wider TMD pores, notably at the -1’, 9’, and 16’ constriction points.

### 2.3 Vestibular Ligand Binding in Open X-ray and Cryo-EM Structures

To verify binding of psychostimulant derivatives to the open state of sTeLIC, we then took advantage of the anomalous X-ray diffraction signal available from the bromine moiety, solving a co-crystal structure of the complex with BrAmp. We co-crystallized the drug with the receptor according to previously established protocols in detergent (n-dodecyl-b-maltoside (DDM) and nonyl-β-d-glucose (NDG)) at pH 8.0 [15]. The resulting crystals diffracted up to 3.2 Å resolution, enabling structure determination by molecular replacement using a previously solved sTeLIC X-ray structure (PDB ID 6FL9 [15]) as a search model (Supplementary Table S2). Anomalous signal from the bromine atom of BrAmp allowed definitive placement in the vestibular pocket of each subunit, with the bromine buried deep in the intrasubunit cavity (Figures 2A and 2B). We obtained a comparable structure with BrPEA to 3.0 Å resolution, verifying equivalent binding poses of the potentiators (Supplementary Figure S1 and Supplementary Table S2). Both structures were otherwise superimposable with previously reported open states of sTeLIC (Cβ RMSD *<* 0.5 Å versus PDB ID 6FLI), with a pore radius (≥ 6 Å) wider than that of open ELIC [25] and sufficient to conduct hydrated sodium ions (Figures 2C, 2D and Supplementary Figure S1A). To verify that these structures were not substantially influenced by crystallization, we also collected cryo-EM data for sTeLIC in detergent with BrAmp. Although the ligand could not be definitively built without the benefit of anomalous diffraction, the protein density corresponded closely to the open X-ray structures (Supplementary Table S1 and Supplementary Figure S2).

We also attempted to obtain an open-state apo structure, that is, without modulatory ligands. Nanodisc samples behaved poorly in our hands under the elevated-pH conditions predicted to activate the channel; hence, supported by its evident insensitivity to membrane mimetic (described above), we prepared sTeLIC grids in detergent at pH 9.0. The resulting cryo-EM structure resolved to 2.6 Å (Supplementary Table S1 and Supplementary Figure S2), and was superimposable with other open states (Cβ RMSD *<* 0.6 Å versus PDB ID 6FLI). In contrast to the closed state, local resolution in all open cryo-EM structures was relatively high in the ECD, and lowest at the inward-facing end of the TMD (Supplementary Figure S2). Even in the absence of known ligands, the vestibular cavities again contained nonprotein densities, which could reasonably accommodate the DDM detergent used for protein solubilization (Supplementary Figure S1D). A previously reported X-ray structure determined without known ligands also contained an unidentified density in this region [15]. Thus, whereas the closed structure contained no apparent density in the extracellular vestibule, it appears that the open-state cavity is prone to binding a variety of hydrophobic compounds, rendering characterization of a genuine apo structure challenging.

### 2.4 Structural Transitions Associated with Alkaline Gating

Comparison of our closed to open structures revealed not only expansion of the TMD pore (Supplementary Figure S4), but also rotation and contraction of the ECD, including the extracellular vestibule. Overall, the ECD contracted by 10 percent (25 Å to Å 23 ECD spread, see Methods for measurement details), and rotated 9 degrees anti-clockwise relative to the TMD (ECD twist). These relatively subtle global motions were associated with notable local changes, including translation at the midpoint of each extracellular b2-b3 loop by 11 Å inward, and for loop C by 5 Å towards the complementary subunit (Figure 3A). At the ECD-TMD interface, loop F also exhibited dramatic remodeling, moving 8 Åtowards the conduction pathway. Interestingly, a similar motion was previously reported for the related bacterial channel DeCLIC, reorganizing an intersubunit site for Ca^2+^ inhibition [27]. In the TMD, the M2-M1(-) distance contracted by 9 Å (21 Å to 12 Å), and the M2–M3 loop moved 6 Å away from the pore (Figure 3A). The closed pore profile (Figure 2C) was similar to those of ELIC [23, 25] and other pLGICs (Supplementary Figure S4A), while the open pore profile was notably wider (Figure 2D and Supplementary Figure S4B).

**Figure 3.**
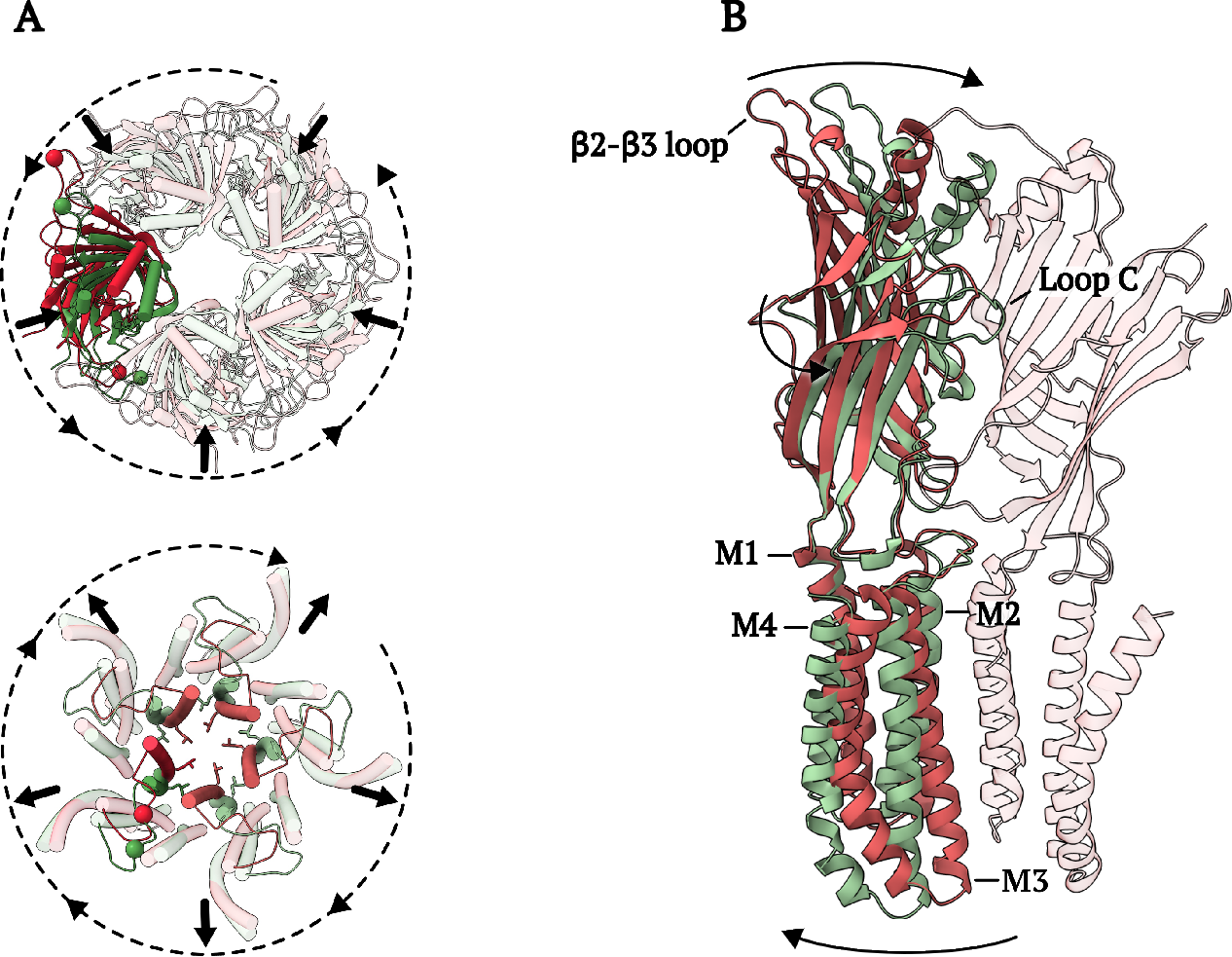
Structural transitions associated with sTeLIC gating. **A)** Extracellular views of superimposed sTeLIC structures in closed (red) and open (green) states, highlighting conformational changes occurring in the ECD (top) and TMD (bottom). A single-subunit ECD and the pore-lining M2-helices are darkened for clarity. Arrows indicate structural rearrangements upon activation, involving ECD contraction, and anticlockwise twist relative to the TMD (top), and TMD expansion (bottom). For reference, the ECD model includes Cβ atoms at the midpoints of the highlighted β2–β3 loop and loop C as spheres (top); the TMD model includes the hydrophobic gate formed by L234 (9’) as sticks, and the midpoint of one M2–M3 loop as spheres (bottom). **B)** Conformational changes in a single sTeLIC subunit, showing closed (red) and open (green) structures aligned on the ECD-TMD interface and viewed from the membrane plane. For reference, the closed-state complementary subunit is shown partially transparent. Arrows indicate ECD contraction and TMD expansion.

### 2.5 Basis for Positive Allosteric Modulation via the ECD Vestibule

In addition to the global ECD motions described above, superposition of a single subunit from closed and open sTeLIC structures revealed local changes in and around the vestibular site (Figure 4A). This site was defined in part by the so-called Ω-loop, a region spanning the b4–b5 strands that has been implicated in subtype-specific modulation [15], as well as the b6 strand on the inner face of the cavity. Whereas the closed state contained no apparent pocket capable of accommodating BrAmp (Figure 4B), all open structures contained such a site, sandwiched between residues including b4-W75 and b6-Y104 (Figure 4C). Specifically, the side chain of W75 occluded the pocket in the closed state, but rotated outward to evacuate it in open structures. Whereas BrAmp could not be docked favorably in the closed state (best pose 17 kcal/mol), it gave rise to favorable energy scores (best pose -4.8 kcal/mol) in the open state, mediated in part by apparent aromatic interactions with Y104. Consistent with these predictions, substituting Ala or Val at either W75 or Y104 significantly reduced BrAmp potentiation of sTeLIC in oocyte recordings (Figure 4E).

**Figure 4.**
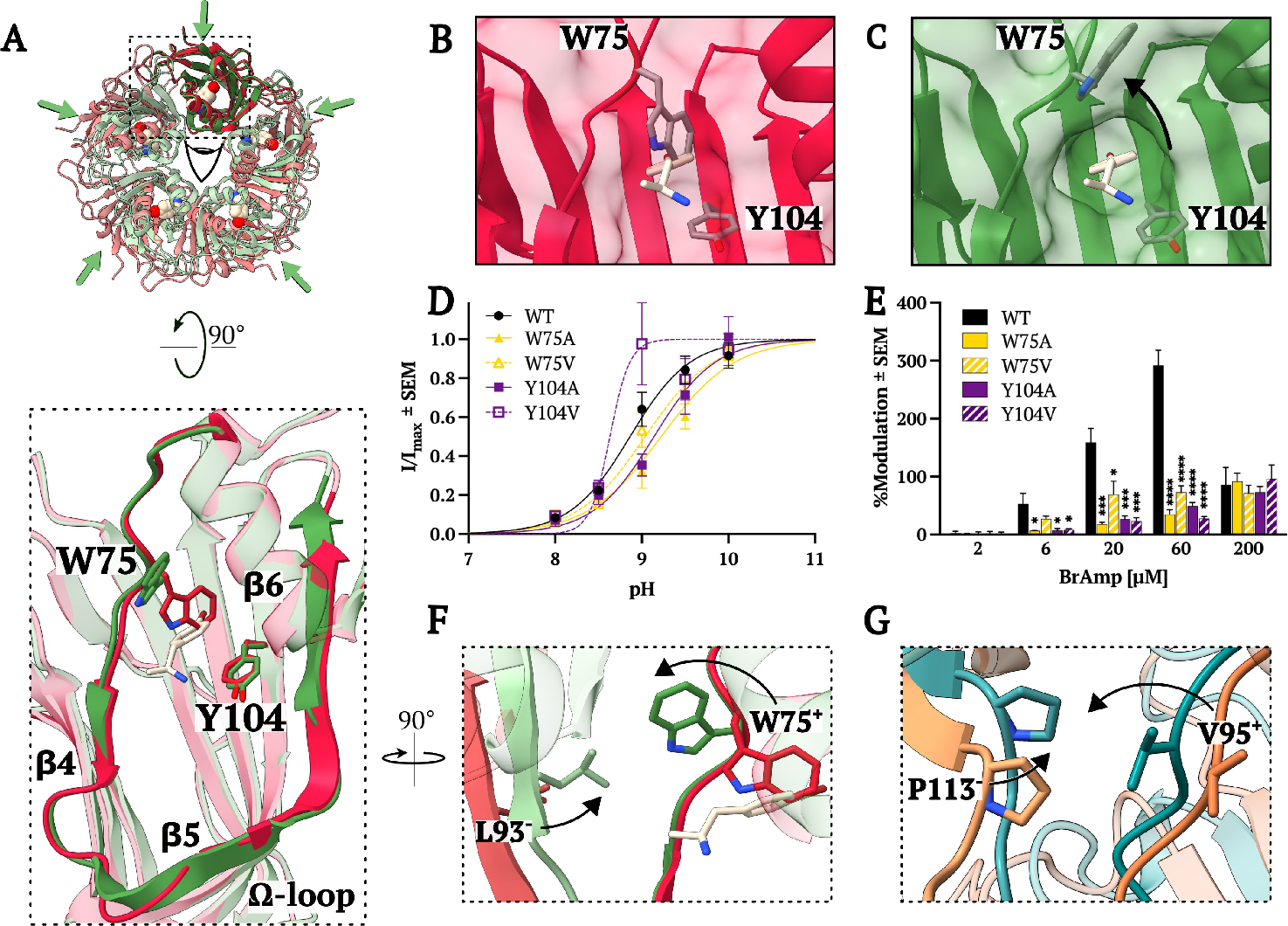
Structural basis for positive allosteric modulation via the ECD vestibule. **A)** Closed (red) and open (with BrAmp, X-ray, green) structures of sTeLIC, viewed fom the extracellular side, showing relative contraction of the ECD upon activation. Inset, superimposing the structures on a single-subunit ECD and viewing from the vestibule (indicated by perspective eye) reveals reorientation of β4-W75 out of the pocket, which is instead occupied by BrAmp (beige) proximal to β 6-Y104 in the open state. **B)** Vestibular pocket in the closed state, viewed as in (A, inset), including a semitransparent molecular surface. BrAmp (beige) from the open X-ray structure is superimposed for perspective. **C)** Vestibular pocket in the open structure (with BrAmp, X-ray), viewed as in (B) including the bound BrAmp (beige) and a semitransparent surface. **D)** pH-response curves for WT and mutant receptors (n *≥* 4) recorded by TEVC electrophysiology in *Xenopus laevis* oocytes. Normalized currents (I/I_*max*_) *±* SEM were fitted by nonlinear regression, yielding EC_20_ values for all constructs around pH 8.5 *±* 0.2. **E)** Percent modulation of sTeLIC current amplitudes upon coactivation of WT and mutant channels at pH 8.5 with BrAmp (n *≥* 3), relative to currents activated without BrAmp immediately before treatment. Significance is relative to WT (*P ≤ 0.05, **P ≤ 0.01, ***P ≤ 0.001, ****P ≤ 0.0001). **F)** Rotated view of superposition as in (A, inset), revealing interactions of β 4-W75 with β 5-L93 on the complementary subunit upon transitioning from closed (red) to open (green). **G)** View as in (F) of a mammalian 5-HT_3*A*_R, showing a cryo-EM structure without BrAmp (PDB ID 6DG8, orange) and a computational model with BrAmp (teal, [28]). Residues V95 and P113, corresponding to sTeLIC W75 and L93 respectively, have been shown to mediate BrAmp modulation, and are shown as sticks.

In a plausible model for state-dependent binding, displacement of the W75 side chain from the vestibular pocket not only facilitates ligand binding, but enables it to make hydrophobic contacts with β5-L93 on the complementary subunit (Figure 4F). In the closed state, an equivalent rotation of W75 would not produce substantial contacts with the neighboring subunit, due to the relative expansion of the ECD. Indeed, multiple substitutions at residue L93 (Ala, Val, Ser, Asp) ablated functional sTeLIC expression in oocytes, consistent with a critical role for this residue in channel gating and/or assembly. In parallel work, we recently reported that mutating equivalent positions in mammalian 5-HT_3*A*_Rs (β4-V95, β6-N125, β5-P113) decreases sensitivity to both 5-HT activation and BrAmp potentiation (Figure 4G) [28]. This putative mechanism may be a specific feature of bacterial pLGICs and 5-HT_3*A*_Rs, in which the vestibular pocket exhibits conserved architecture and intersubunit interactions (Supplementary Figure S5A); in contrast, this cavity in eukaryotic receptors for GABA, glutamate, glycine, and acetylcholine are notably different and largely occluded (Supplementary Figure S5B).

## 3. Discussion

In this work, new cryo-EM and X-ray structures in combination with engineered sitedirected mutations support the presence of a druggable pocket in the sTeLIC ECD vestibule, that binds stimulants in a state-dependent manner (Figure 5). In a plausible mechanistic model, global contraction of the upper ECD and local rearrangement of the W75 side chain facilitate contacts between Ω-loop residues (e.g. β4-W75 and β5-L93), and render the vestibular pocket accessible to ligands (mediated in part by β6-Y104), promoting a potentiated state. The vestibular site appears to accommodate a variety of amphiphilic ligands, including BrAmp, BrPEA, FFC-8, and DDM, all producing superimposable protein conformations despite variations in pH (7.5 or 9.0), membrane mimetic (detergent micelles or lipid nanodiscs), and structure determination method (X-ray crystallography or cryo-EM). In all cases, relative enhancement in local ECD resolution in open versus closed cryo-EM structures (Supplementary Figure S2) suggests that stabilization in this region promotes channel activation, possibly via vestibular contacts. Interestingly, a similar correlation between activation and ECD stabilization has been reported for the homologous prokaryotic channel GLIC [24]; GLIC has also been shown to bind modulators such as dicarboxylates in its vestibular site, though the relative influence of vestibular versus orthosteric binding has been unclear [29].

**Figure 5.**
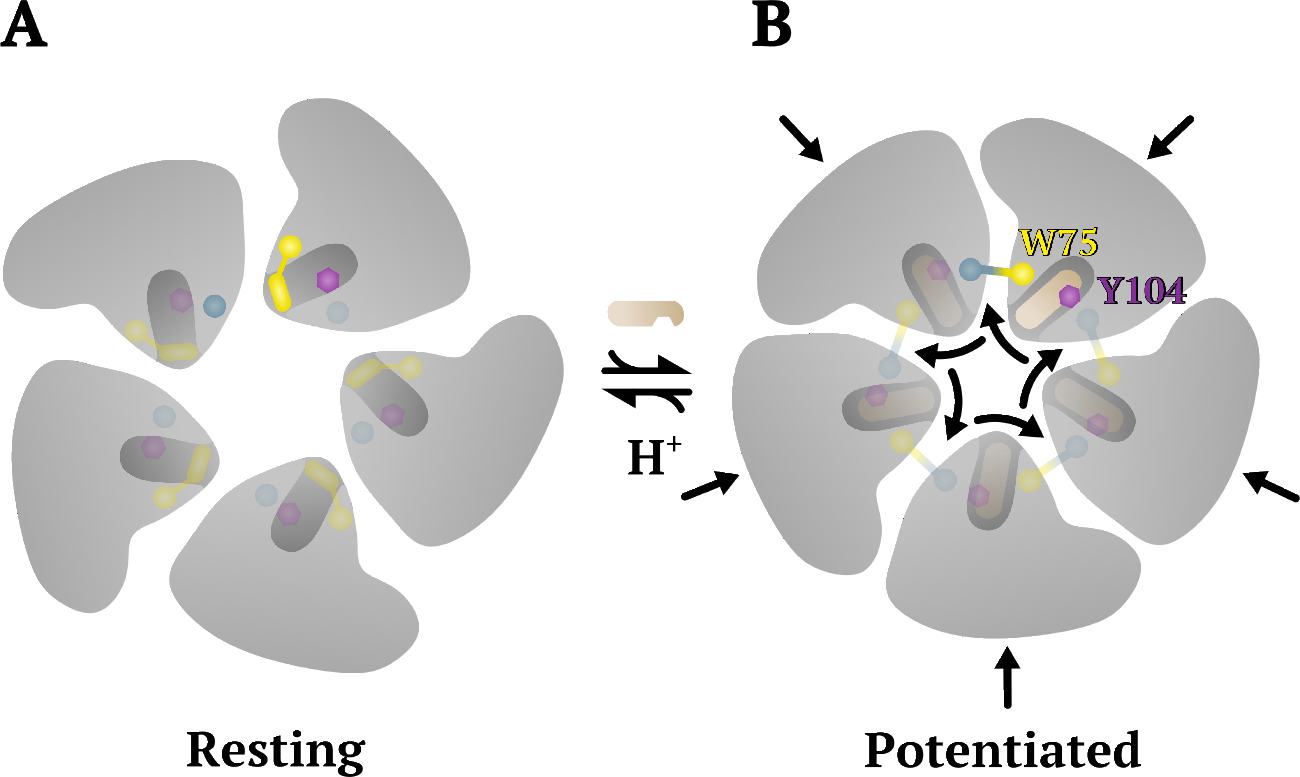
Proposed mechanism of allosteric enhancement via the vestibular site. **A)** Cartoon of the ECD in a resting state, with an expanded pentamer, and W75 (yellow) occluding the vestibular site. **B)** Upon activation by deprotonation and/or modulator binding (reaction arrows), the ECD contracts (straight arrows), bringing the β4 and β5 strands on either end of the Ω-loop in closer proximity to one another. Binding of modulator at β6-Y104 (purple) is enabled by reorientation of β4-W75 (yellow) out from the vestibular site (curved arrows), enabling it to interact directly with β5-L93 (blue) on the complementary subunit and potentially stabilizing the activated state. For clarity, intersubunit contacts associated with the top-right subunit are rendered opaque.

Psychostimulant derivatives including BrAmp, BrPEA, and PEA act as positive vestibular modulators of sTeLIC, generally increasing in the low-to-high micromolar range. A modest decrease in potency at the highest BrAmp and BrPEA concentrations tested (200 µM and 1 mM, respectively) may be related to confounding effects of desensitization, ligand solubility, and/or alternative binding sites. Both of the brominated compounds were more potent than PEA, indicating that bromine interactions deep in the vestibular pocket promote binding; interestingly, controlled substances including 4-bromo-2,5-dimethoxyphenethylamine (2C-B) contain a bromine atom at an equivalent aromatic position [30]. Notably, several diverse ligands appear capable of binding and modulating this site in sTeLIC and its relatives, including FFC-8, dicarboxylates (in GLIC) [14], and flurazepam (in ELIC) [10].

No structures of eukaryotic pLGICs have been captured thus far with ligands in the vestibular site. Recent work by Brams and colleagues indicates this pocket to be occluded in several eukaryotic pLGICs, but selectively accessible to ligands in 5-HT_3*A*_Rs [19]. Their work further demonstrates that covalent labeling in the vestibular site modifies 5-HT_3*A*_R currents, supporting a functional role in receptor modulation [19]. In parallel with the present study, we have observed BrAmp in particular to potentiate 5-HT_3*A*_Rs even more potently than sTeLIC [28]. Furthermore, BrAmp modulation of 5-HT_3*A*_Rs is sensitive to mutations at positions equivalent to sTeLIC β4-W75, β5-L93, and β6-Y104, and appears to require a similar rearrangement of the β4 residue out of the vestibular site and into contact with the complementary β5 [28]. Taken together, these findings support a general mechanism of pocket opening and ligand modulation in a subset of pLGICs including mammalian 5-HT_3*A*_Rs.

As a model system for biomedically relevant pLGICs, our structures of sTeLIC offer direct visualization of vestibular modulation by stimulant derivatives. In animals, amphetamines have long been known to increase serotonin levels by inhibiting presynaptic monoamine transporters and monoamine oxidases, enzymes responsible for serotonin breakdown [31–34]. However, psychostimulants can induce a range of side effects of unclear origin, including nausea, hypomania, and restlessness [35–37]. Indeed, 5-HT_3*A*_R inhibitors include important treatments for chemotherapy-induced nausea; in contrast, stimulant potentiation could underlie undesirable gastrointestinal effects. Thus, our structure-function data offer a mechanistic basis to target or tailor pharmacological tools against a relatively unexplored site, and to understand the complex allosteric gating landscape of membrane receptors.

## 4. Materials and Methods

### 4.1 Mutagenesis Design and Generation

We used molecular visualization and docking tools in UCSF Chimera 1.16 [38] to identify residues in the vestibular binding site in closed and open structures of sTeLIC. BrAmp was prepared for docking using the Dock Prep [39] tool with default settings, which adds polar hydrogens and charges. Local docking simulations were then conducted using the AutoDock Vina [40] module, with a 15 Å × 13 Å × 19 Å search volume enclosing and centered in the vestibular pocket of the closed cryo-EM structure or an open X-ray structure (PDB ID 6FL9 [15]).

Mutations were generated in sTeLIC inserted in the vector pMT3, using the GeneArt Site-Directed Mutagenesis System (Thermo Fisher Scientific) and commercial mutagenic primers (IDT). Mutant cDNAs were then verified by Sanger sequencing (Eurofins Genomics) and purified on larger scale using a HiSpeed Plasmid Midi Kit (Qiagen).

### 4.2 Oocyte Preparation

Oocytes from female *Xenopus laevis* (EcoCyte Bioscience) were injected via the animal pole with 6 ng/32.2 nl sTeLIC WT or mutant cDNA using a Nanoject II microinjector (Drummond Scientific). After injection, oocytes were kept in a 48-well plate containing modified Barths solution (88 mM NaCl, 1 mM KCl, 2.4 mM NaHCO_3_, 0.91 mM CaCl_2_, 0.82 mM MgSO_4_, 0.33 mM Ca(NO_3_)_2_, 10 mM HEPES, 0.5 mM theophylline, 0.1 mM G418, 17 mM streptomycin, 10,000 U/l penicillin and 2 mM sodium pyruvate, adjusted to pH 7.5) at 13°C.

### 4.3 Oocyte Electrophysiology

BrAmp-modulated currents were recorded 5–12 days after cDNA injection using a two-electrode voltage-clamp. The oocyte membrane was voltage-clamped using two recording needles (5 - 50 MΩ) filled with 3 M KCl and two electrodes connecting ground stages and the bath chamber through 3 M KCl-agar bridges. Oocytes were perfused with the running buffer (123 mM NaCl, 2 mM KCl and 2 mM MgSO_4_, and adjusted to pH 7.5) at a flow rate of roughly 1 mL/min. Oocytes were clamped at -40 mV using an OC-725C voltage clamp (Warner Instruments). A Digidata 1440A (Molecular Devices) was used to sample and digitize data using Clampex (Axon Instruments). Currents were filtered at a frequency of 1 kHz and analyzed using Clampfit (Axon Instruments).

Before each recording occasion, we freshly prepared a 1-M stock solution of modulator (BrAmp, BrPEA, PEA, or FFC-8) in DMSO. Solutions with varying modulator concentrations were then prepared in a buffer pH-adjusted to produce 20 percent maximal current (pH 8.5 for all variants). pH-activation buffer, in the absence or presence of modulator, was applied for 30 sec, followed by 7 min washout. After each modulator application, the two following applications were solely ∼ EC_20_ buffer, to verify a return to baseline behavior.

### 4.4 Statistics

All statistical analyses were carried out in Prism 9.5.0 (GraphPad Software). Data are represented as mean *±* SEM and analyzed with unpaired t-tests, with significant effects set at P *<* 0.05. Proton activation curves were calculated using the equation, 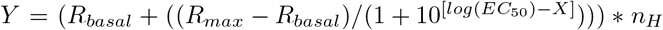, where Y is the activation response, R_*max*_ is the maximal response, R_*basal*_ is the baseline, X is the proton concentration, EC_50_ is the concentration of a drug that induces 50 % of maximal response, and n_*H*_ is the Hill coefficient. Percent BrAmp modulation was calculated as %*modulation* = ((*R*_*A*_ *-R*_0_)*/R*_0_) *\** 100, where R_*A*_ represents activation responses, and R_0_ portrays response occurring right before the R_*A*_ response.

### 4.5 sTeLIC Expression and Purification

sTeLIC was expressed and purified as described previously [15]. Briefly, C43(DE3) *Escherichia coli* cells were transformed with vector pET20b containing maltose-binding protein (MBP) fused to the N-terminus of sTeLIC. Cells were inoculated 1:100 into 2xYT media with 100 µg/ml ampicillin, grown at 37°C to OD600 = 0.8, induced with 0.1 - 0.4 mM isopropyl-β-D-1-thiogalactopyranoside (IPTG), and shaken overnight at 20°C. The overnight culture was harvested and resuspended in buffer A (20 mM Tris pH 8.0, 300 mM NaCl) supplemented with EDTA-free protease inhibitor (Thermo Fisher Scientific) and either benzonase nuclease or lysozyme plus DNase I. The cell membranes were solubilized in a 4 % DDM buffer at 4°C for 3–16 hours. After membrane treatment, the extracted recombinant protein was purified on amylose resin, eluting with buffer B containing 20 - 50 mM D-(+)-maltose monohydrate, and further purified by size exclusion on a Superose TM6 Increase 10/300 GL column. MBP was subsequently separated from sTeLIC by thrombin cleavage and a second round of gel filtration in a buffer containing 20 mM Tris pH 8.0, 300 mM NaCl and 0.02 % DDM. Purified protein was pooled and concentrated to 12 mg/mL.

The plasmid for SapA expression was a gift from Salipro Biotech AB, and the purification of SapA followed previously published protocols [41]. Saposin nanodisc reconstitution started with mixing sTeLIC, SapA and polar brain lipid at molar ratio 1:15:150, then incubating the mixture on ice for 1 h. Bio-Beads SM-2 (Bio-Rad) were added to the mixture, then gently shaken overnight at 4°C. The next day, the supernatant was collected and purified in 20 mM HEPES pH 7.5, 100 mM NaCl, by size-exclusion chromatography on a Superose TM6 Increase 10/300 GL column (Cytiva). Peak fractions were collected and concentrated to *∼* 3 mg/mL.

### 4.6 Crystallization, Data Processing and Structure Determination

In order to produce large, well diffracting crystals, the purified protein was first mixed with 65 mM NDG at 4:1 volume ratio and incubated on ice for 20 min. Co-crystallization of sTeLIC with BrAmp or BrPEA was performed by mixing the NDG-protein mixture with the desired molecule solubilized in 100 % DMSO at a final concentration of 3 mM. Crystals were then grown by mixing the obtained protein solution with an equal volume of reservoir solution containing 100 mM Tris pH 8.0, 150 mM MgCl_2_, 3 % DMSO, and 30-35 % PEG 200 at 1:1 volume ratio. Mixtures were equilibrated against a 1 mL reservoir solution using the hanging drop method at 18°C. Co-crystals appeared within two weeks and grew to full size within a month. Prior to data collection crystals were mounted on nylon loops and flash frozen in liquid nitrogen, using the crystallization solution as a cryo-protectant.

Diffraction data sets were collected at synchrotron Soleil (PROXIMA-1 and PROXIMA-2A) and processed using XDS [42] and STARANISO (Global Phasing Limited) softwares. The structure was solved by molecular replacement using a previously solved sTeLIC X-ray structure (PDB ID 6FL9) as a search model by Phaser-MR in CCP4 [43]. Refinement was performed using REFMAC5 software from the CCP4 suite with non-crystallographic symmetry (NCS) restraints applied. Anomalous maps were calculated using CAD software from the CCP4 suite and averaged in Coot [44].

### 4.7 Cryo-EM Sample Preparation and Data Acquisition

For saposin-nanodisc grids, 0.3 µL FFC-8 (Anatrace) or CHAPS (Anatrace) stock was added to 2.7 µL sample immediately before grid freezing, to a final concentration of 2 mM. The mixtures were applied to glow-discharged R1.2/1.3 300 mesh Au grids (Quantifoil), blotted with force 0 for 2 s, and plunged into liquid ethane using a Vitrobot Mark IV (Thermo Fisher Scientific) at 20°C. For the detergent sample at pH 7.5, 100 µM BrAmp was added to *∼* 3 mg/ml protein. 3 µL of the mixture was then applied to glow-discharged 1.2/1.3 300 mesh Cu grids (Quantifoil). For the detergent sample at pH 9, 3 µL of a *∼* 7 mg/ml sample was directly applied to 1.2/1.3 300 mesh Au grids (Ultrafoil) without any additives. For both detergent samples, the grids were then blotted with force 0 for 2 s and plunged into liquid ethane using a Vitrobot Mark IV (Thermo Fisher Scientific) at 4°C and 100 % humidity. Cryo-EM data were collected on a Titan Krios microscope (Thermo Fisher Scientific) operated at 300 kV with a K3 Summit detector (Gatan), using EPU automated collection software (Thermo Fisher Scientific). The total dose in each nanodisc dataset was *∼* 45 electrons/Å^2^ and the defocus range set from -0.8 to -2.4 µm. The total dose in the detergent datasets were *∼* 40 electrons/Å ^2^ (pH 7 with BrAmp) or *∼* 36 electrons/Å ^2^ (pH 9), and the defocus range set from -0.8 to -3 µm.

### 4.8 Image Processing

Dose-fractionated images in super-resolution mode were internally gain-normalized and binned by 2 in EPU (Thermo Fisher Scientific) during data collection. Motion correction of the dose-fractionated images was done in RELION 4.0 [45]. Contrast transfer functions (CTFs) were estimated on the motion-corrected images using CTFFIND4.1 [46] and particles were automatically picked with 2D references generated from manually picked particles (detergent structures), or with Topaz 0.2.5 [47] using the internal trained model (nanodisc structures). The rest of the processing was performed in RELION 4.0. First the particles were binned by 4 and extracted. 2D classification was run for the binned particles, then the classes with recognizable protein features were selected and re-extracted binned by 2. For the detergent structures, the initial model was generated using ab initio reconstruction. For the nanodisc structures, the initial model was generated from the PDB of a previously published sTeLIC crystal structure (PDB ID 6FL9) using the molmap command in UCSF Chimera [38]. 3D classification was used to check the structural heterogeneity and further clean the dataset. High quality classes were selected and re-extracted without binning, and these particles were then used for refinement with C5 symmetry imposed, followed by multiple rounds of CtfRefine and polishing to improve resolution.

### 4.9 Model Building

A previously published crystal structure (PDB ID 6FL9) was used as the starting model. The structure was first rigid body fitted into each cryo-EM map in ChimeraX [48], then flexible fitted by ISOLDE [49]. After that, each model was manually adjusted, and ligands and lipids added in Coot 0.9.5 [44]. Models were optimized with PHENIX real-space refinement [50] and validated by MolProbity [51].

### 4.10 Gating Analysis

ECD twist, ECD spread and M2-M1(-) distance were calculated based on a previously published script from [52]. Briefly, ECD twist is defined by the dihedral angle between a) the Ca-center of mass (COM) of a single-subunit ECD, b) the Cβ-COM of the full ECD, c) the Cβ-COM of the full TMD, and d) the Cβ-COM of a single-subunit TMD. ECD spread is defined by the distance from the Cβ-COM of the full ECD to the Cβ-COM of a single-subunit ECD, and M2-M1(-) distance by the distance between the COM of a full M2-helix and that of the full complementary M1-helix. Specific loop movements are based on shifts in the Cα atoms of P51 (for the β2–β3 loop), F177 (for loop C), L157 (for loop F), or L250 (for the M2–M3 loop) between aligned closed and open states.

## Acknowledgments

E.K. and O.A. were supported by PhD fellowships from Sven och Lilly Lawskis fond. C.F. was supported by grant FV-5.1.2-0523-19 from Stockholm University, and R.J.H. and E.L. by grants from the Swedish Research Council (2019-02433, 2021-05806) and Swedish e-Science Research Centre. We also thank the staff of synchrotron Soleil, and especially Pierre Legrand, for help in data collection. Cryo-EM data were collected at the Cryo-EM Swedish National Facility funded by the Knut and Alice Wallenberg, Family Erling Persson and Kempe Foundations, SciLifeLab and Stockholm University.

## Data availability

Cryo-EM densities, including sharpened and unsharpened maps, both half-maps, and the mask used for final FSC calculation, have been deposited in the EMDB at accession numbers EMD-50031 (closed), EMD-50030 (open FFC-8), EMD-50195 (open BrAmp), and EMD-50194 (open pH 9). Coordinates of cryo-EM and X-ray models have been deposited in the PDB at accession numbers 9EX6 (closed), 9EWL (open BrAmp, X-ray), 9EWA (open BrPEA, X-ray), 9F5O (open BrAmp, cryo-EM), 9EX4 (open FFC-8, cryo-EM), and 9F5N (open pH 9, cryo-EM).

## Author contributions

Conceptualization: E.K., R.J.H., M.D.; electrophysiology: E.K.; cryo-EM structure determination and figure preparation: O.A., C.F.; X-ray structure determination and figure preparation: Z.F., A.H.; model building support: Y.Z.; writing - original draft: E.K., M.D., C.F., R.J.H.; writing - review and editing: E.K., O.A., C.F., R.J.H., M.D., E.L.; funding acquisition: E.L., M.D.

## Abbreviations

5-HT_3*A*_R: Serotonin-3A receptor
BrAmp: 4-Bromoamphetamine
BrCA: 4-Bromocinnamate
BrPEA: 4-Bromophenethylamine
Cryo-EM: Cryogenic electron microscopy
EC_50_: Half-maximal effective concentration
ECD: Extracellular domain
FFC-8: Fluorinated fos-choline-8
ICD: Intracellular domain
pLGIC: Pentameric ligand-gated ion channel
SEM: Standard error of mean
TEVC: Two-electrode voltage-clamp
TMD: Transmembrane domain
WT: Wild type

