## Supplementary Information for "Vestibular Modulation by Stimulant Derivatives in a Pentameric Ligand-Gated Ion Channel"

### 5 Supplementary Information

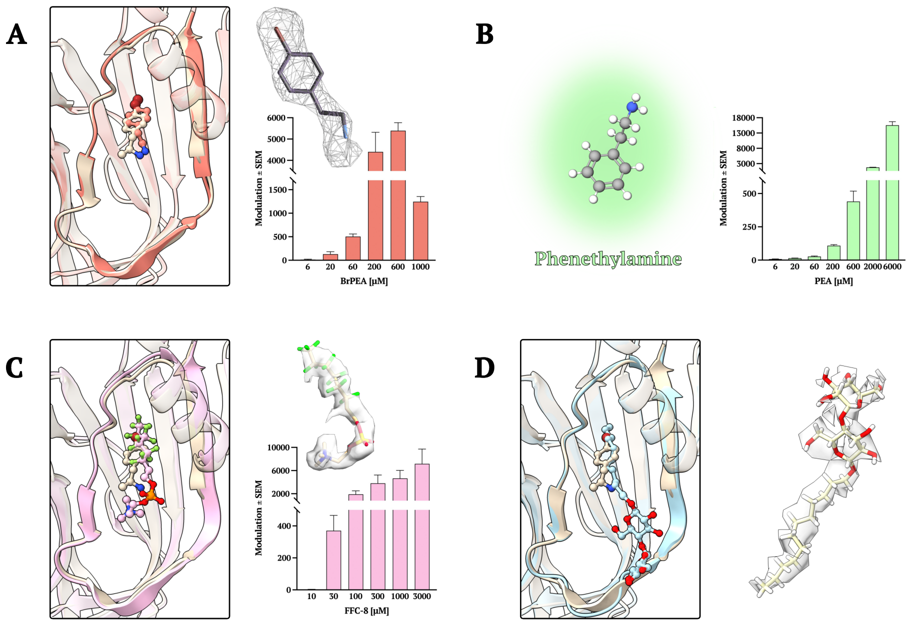


**Figure S1. Other experimentally determined positive allosteric modulators operating at the ECD vestibular site. A)** BrPEA (coral), **C)** FFC-8 (pink), and **D)** DDM (light blue) were superimposed on top of the BrAmp (beige) structure, along with column graphs and density maps for each molecule (n $\geq$ 4). **B)** The column graph and ball-and-stick representation for PEA (light green).

**Table S1 Cryo-EM data collection, refinement and validation statistics.**

| | Closed (9EX6) | Open, FFC-8 (9EX4) | Open, pH 9 (9F5N) | Open, BrAmp (9F5O) |
|---|---|---|---|---|
| | Nanodisc | | Detergent | |
| **Data Processing** | | | | |
| **Magnification** | 130.000 | 130.000 | 130.000 | 130.000 |
| **Voltage (kV)** | 300 | 300 | 300 | 300 |
| **Electron Exposure ($e^-$/Å$^2$)** | 48.38 | 43.44 | 36.6 | 40.27 |
| **Defocus Range (µm)** | -0.8 to -1.8 | -0.8 to -1.8 | -0.8 to -3 | -0.8 to -3 |
| **Pixel Size (Å)** | 0.6725 | 0.6725 | 0.65 | 0.65 |
| **Symmetry Imposed** | C5 | C5 | C5 | C5 |
| **Final Particles** | 136.209 | 148.478 | 218 875 | 77 517 |
| **Map Resolution (Å)** | 2.35 | 2.19 | 2.56 | 4.16 |
| **FSC Threshold** | 0.143 | 0.143 | 0.143 | 0.143 |
| **Refinement** | | | | |
| **Map Sharpening B Factor (Å$^2$)** | -62.3 | -53.5 | -90.8 | -296.49 |
| **Non-Hydrogen Atoms** | 13225 | 13250 | 12500 | 11630 |
| **Protein Residues** | 1550 | 1550 | 1550 | 1550 |
| **Ligands** | 35 | 20 | 5 | 0 |
| **B factors (Å$^2$)** | | | | |
| Protein | 40.60 | 27.71 | 33.39 | 41.81 |
| Ligand | 42.09 | 24.23 | 39.48 | - |
| **RMS Deviations** | | | | |
| Bond Lengths (Å) | 0.007 | 0.002 | 0.007 | 0.005 |
| Bond Angles (°) | 1.143 | 0.386 | 0.733 | 0.675 |
| **Validation** | | | | |
| MolProbity Score | 1.31 | 1.19 | 1.34 | 1.71 |
| Clashscore | 3.15 | 4.02 | 5.19 | 6.77 |
| Poor Rotamers (%) | 0.80 | 0.00 | 0.00 | 0.00 |
| **Ramachandran Plot** | | | | |
| Favored (%) | 96.75 | 99.03 | 97.73 | 95.13 |
| Allowed (%) | 3.25 | 0.97 | 2.27 | 4.87 |
| Outliers (%) | 0.00 | 0.00 | 0.00 | 0.00 |

**Table S2 X-ray data collection and refinement statistics.** Statistics for the highest-resolution shell are shown in parentheses.

| | Open, BrAmp (9EWL) | Open, BrPEA (9EWA) |
|---|---|---|
| **Data Processing** | | |
| **Wavelength** | 0.919 | 0.919 |
| **Resolution Range** | 24.88 to 3.2 (3.314 to 3.2) | 25 to 3.01 (3.117 to 3.01) |
| **Space Group** | C121 | I121 |
| **Unit Cell** | 218.38 114.32 142.44 90 113.052 90 | 143.21 112.345 210.796 90 106.178 90 |
| **Total Reflections** | 66373 (2032) | 126885 (12481) |
| **Unique Reflections** | 33187 (1016) | 63534 (6069) |
| **Multiplicity** | 2.0 (2.0) | 2.0 (2.0) |
| **Completeness (%)** | 62.05 (19.05) | 99.03 (96.04) |
| **Mean I/sigma(I)** | 15.11 (3.59) | 6.94 (1.08) |
| **Wilson B-factor** | 81.99 | 82.35 |
| **R-merge** | 0.1033 (0.3322) | 0.05867 (0.4796) |
| **CC1/2** | 0.98 (0.724) | 0.997 (0.891) |
| **CC*** | 0.995 (0.916) | 0.999 (0.971) |
| **Refinement** | | |
| **R-work** | 0.2244 (0.3059) | 0.2731 (0.4742) |
| **R-free** | 0.2561 (0.3100) | 0.3093 (0.4481) |
| **Non-Hydrogen Atoms** | 13005 | 13062 |
| Macromolecules | 12830 | 12848 |
| Ligands | 139 | 155 |
| Solvent | 36 | 59 |
| **Protein Residues** | 1550 | 1552 |
| **B factors (Å$^2$)** | | |
| Macromolecules | 81.14 | 105.30 |
| Ligands | 122.08 | 125.99 |
| Solvent | 47.23 | 65.71 |
| **RMS Deviations** | | |
| Bond Lengths (Å) | 0.013 | 0.013 |
| Bond Angles (°) | 1.83 | 1.76 |
| **Ramachandran Plot** | | |
| Favored (%) | 94.16 | 95.14 |
| Allowed (%) | 5.84 | 4.80 |
| Outliers (%) | 0.00 | 0.06 |

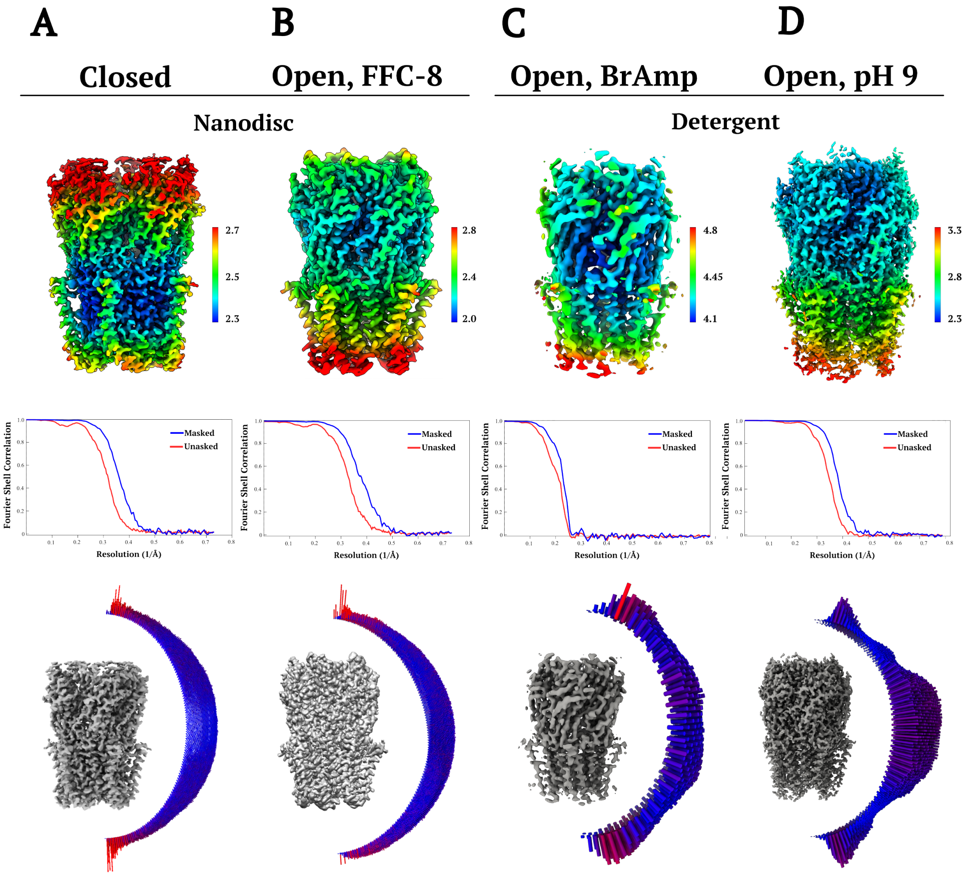


**Figure S2. Local resolution and angular distributions for sTeLIC cryo-EM reconstructions in the absence and presence of various modulators.** Top, Coulombic density colored by local resolution according to scalebar at right. Middle, Fourier shell correlation (FSC) curves before (red) and after (blue) the application of a soft mask in RELION 4.0 [45]. Bottom, angular distribution of particles in the final map. The height is proportional to the number of particles assigned to each view, colored blue-to-red for least-to-most populated views). Reconstructions are shown for **A)** closed and **B)** open (with FFC-8) states in nanodiscs, and open states (at **C)** pH 7.5 with BrAmp, and **D)** pH 9) in detergent.

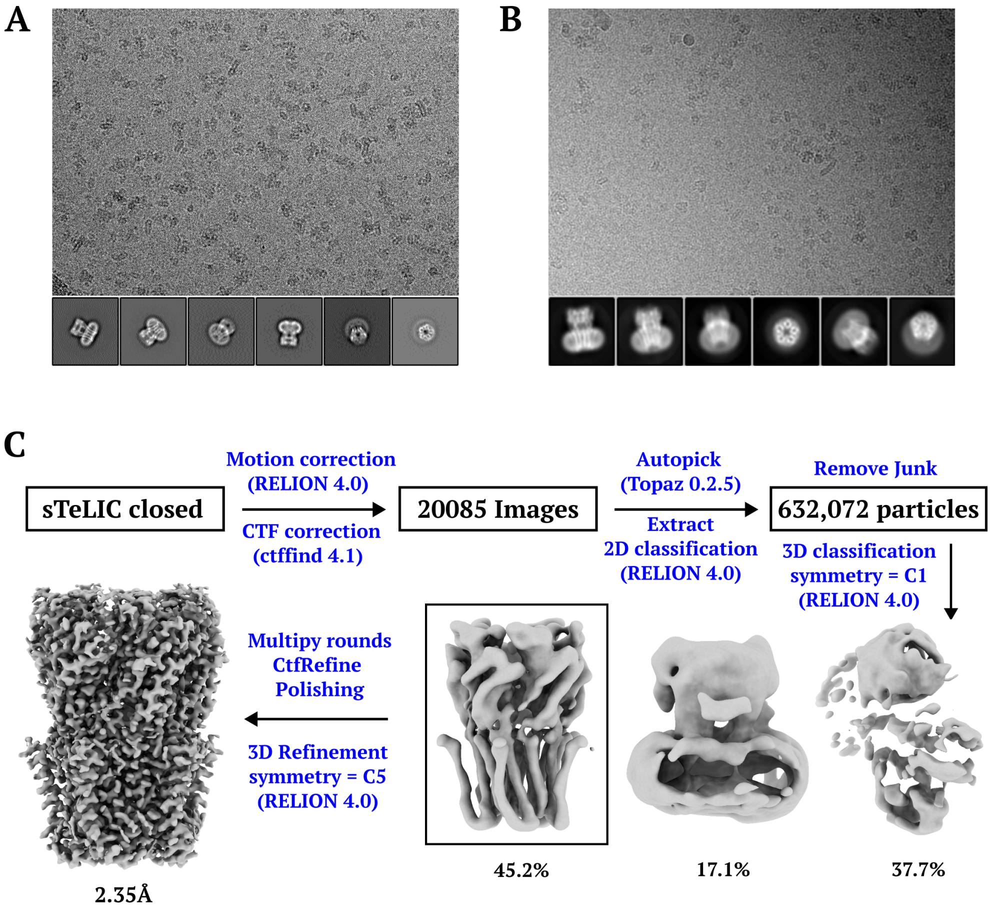


**Figure S3. Cryo-EM data processing workflow.** Representative micrographs of sTeLIC in vitreous ice **A)** in nanodiscs at pH 7.5 and **B)** in DDM at pH 9. Inset, selected 2D classes showing various orientations. **C)** Processing pipeline for a representative sTeLIC reconstruction.

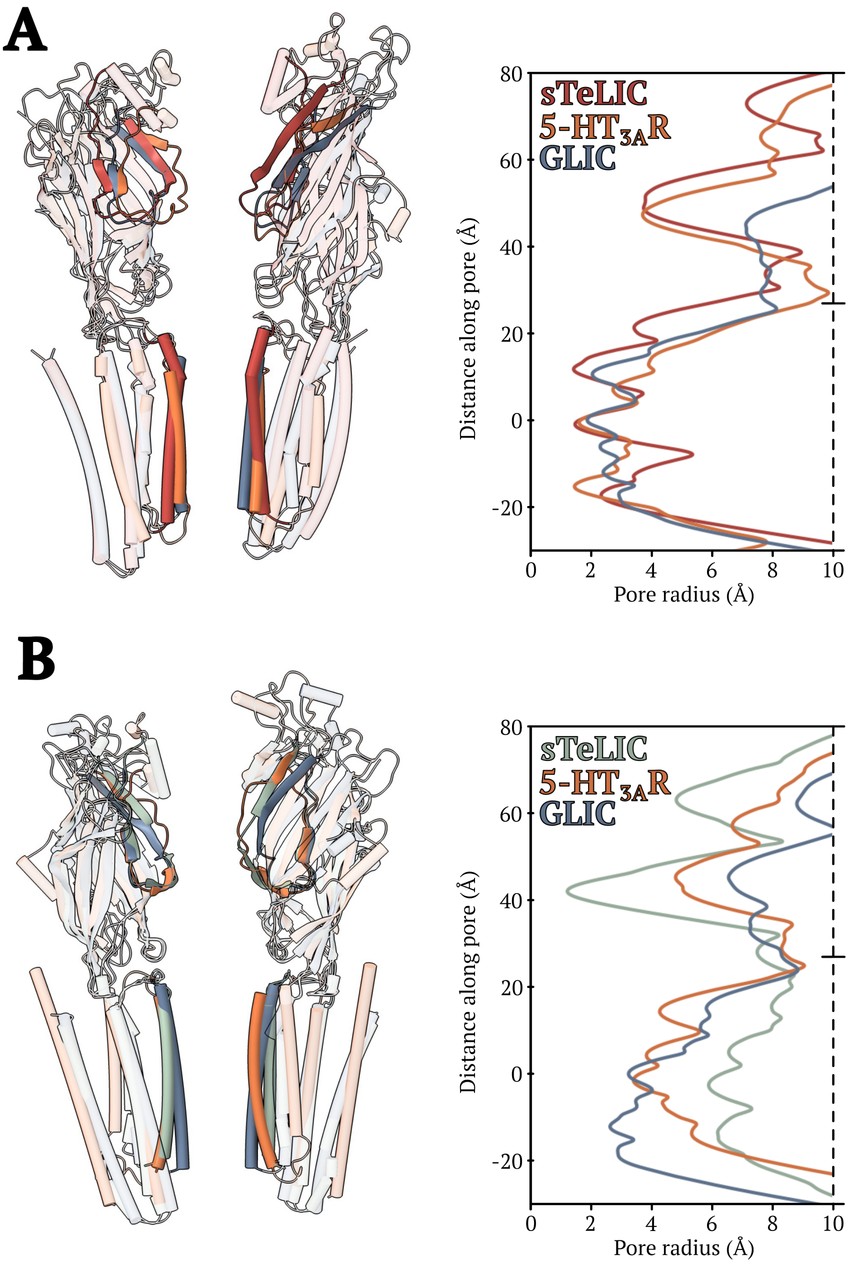


**Figure S4. Structural comparison of sTeLIC to comparable states of prokaryotic and eukaryotic pLGICs. A)** Three pLGICs in presumed resting states, including sTeLIC (red), a 5-$HT_{3A}$R (PDB ID 6BE1, orange), and GLIC (PDB ID 4NPQ, blue), viewed from the membrane plane. For clarity, only two opposing subunits of each channel are shown, with highlighted Ω-loops and M2-helices. Inset, pore profiles. **B)** Three pLGICs in presumed open states, including sTeLIC (X-ray structure with BrAmp, green), a 5-$HT_{3A}$R (PDB ID 6DG8, orange), and GLIC (PDB ID 4HFI, blue), otherwise viewed as in (A).

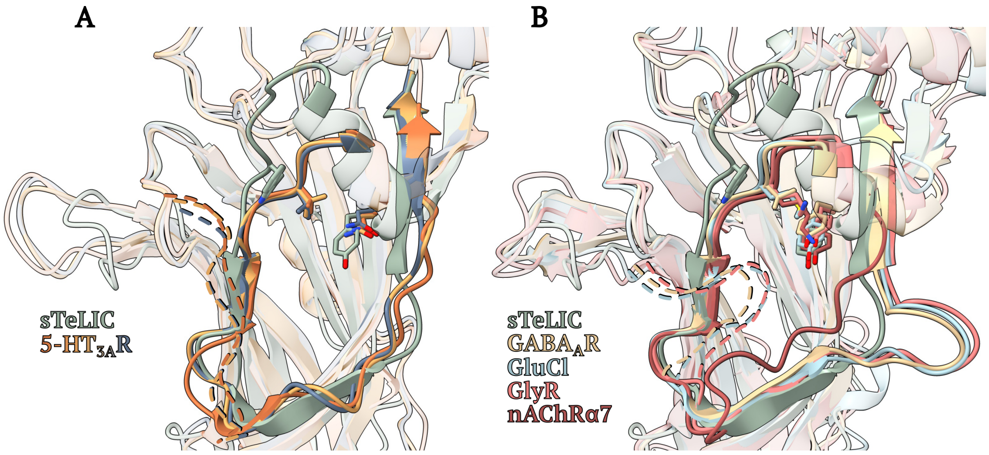


**Figure S5. Structural alignment of a single sTeLIC vestibular pocket with various activated pLGICs. A)** Superimposed sTeLIC (X-ray structure with BrAmp, green) with three active 5-HT$_{3A}$R structures (PDB ID 6HIN, blue; PDB ID 6DG8, orange; PDB ID 6Y5A, yellow), aligned on a single-subunit ECD. Residues implicated in BrAmp modulation of sTeLIC, and at corresponding positions in other pLGICs, are depicted as sticks. The open conformation of the Ω-loop creates an accessible binding site, and brings the β4 strand proximal to β5 of the complementary subunit (dashed ribbons). **B)** Superimposed sTeLIC (X-ray structure with BrAmp, green) with four types of eukaryotic pLGICs. In the α7 nicotinic acetylcholine receptor (nAChRα7, PDB ID 7KOX, red), the Ω-loop collapses into the vestibular pocket of the same subunit. In the GABA$_A$ receptor (α1 subunit, PDB ID 6X3Z, light yellow), GluCl (PDB ID 3RIF, light blue) and glycine receptor (PDB ID 6PM6, pink), the Ω-loop from the complementary subunit (dashed ribbons) extends across the interface to occlude the pocket.
